# Ultra-High Field ^31^P functional Magnetic Resonance Spectroscopy Reveals NAD^+^ Dynamics in Brain Energy Metabolism during Visual Stimulation

**DOI:** 10.1101/2025.11.02.686085

**Authors:** Antonia Kaiser, Fatemeh Anvari Vind, Joao M.N. Duarte, Ileana Jelescu, Yan Lin, Xin Yu, Mark Widmaier, Daniel Wenz, Lijing Xin

**Affiliations:** CIBM Center for Biomedical Imaging, École Polytechnique Fédérale de Lausanne (EPFL), Switzerland; Laboratory for Functional and Metabolic Imaging, École Polytechnique Fédérale de Lausanne (EPFL), Lausanne, Switzerland; Department of Radiology, Lausanne University Hospital and School of Biology and Medicine, University of Lausanne, Lausanne, Switzerland; Radiology Department, Second Affiliated Hospital, Shantou University Medical College, China; Department of Biomedical Engineering, Case Western Reserve University, Cleveland, Ohio, United States; Department of Experimental Medical Science (EMV), Faculty of Medicine, Lund University, Lund, Sweden; Wallenberg Centre for Molecular Medicine, Lund University, Lund, Sweden

**Author notes:** Corresponding author: Lijing Xin, Ph.D., EPFL AVP CP CIBM-AIT, Station 6, CH-1015 Lausanne, Switzerland.

**Keywords:** ^31^P, fMRS, energy metabolism, NAD^+^(H), visual stimulation

## Abstract

We investigated dynamic changes in nicotinamide adenine dinucleotide (NAD□) metabolism in the human occipital lobe using ultra-high field ^31^P functional magnetic resonance spectroscopy (fMRS) at 7 Tesla. Twenty-five healthy volunteers (mean age 24 ± 4 years, 10 female) performed a visual task alternating between fixation and flashing checkerboard stimuli. ^31^P MRS spectra were acquired from a visual cortex voxel functionally localized by prior fMRI. Linear mixed-effects modelling revealed a significant reduction in NAD□ concentrations during the first stimulation block, while no significant change was observed during the second block. No significant changes were observed for other high-energy phosphate metabolites (ATP, phosphocreatine, and inorganic phosphate), indicating specificity in the NAD□ response. Exploratory analyses, dividing the blocks in two halves, suggested further reductions in NAD□ and tNAD in the second halves of both stimulation blocks, though these trends were not statistically significant. Our findings demonstrate the feasibility of using fMRS at 7T to detect stimulus-induced dynamics in cerebral NAD□ metabolism *in vivo*, providing insights into the interplay between glycolysis and oxidative phosphorylation during neural activation.

## INTRODUCTION

Nicotinamide adenine dinucleotide (NAD^+^) and its reduced form NAD + hydrogen (NADH) constitute a pivotal reduction-oxidation (redox) coenzyme pair that is integral to cellular bioenergetics. Perturbations of the NAD^+^/NADH balance have been reported in aging, neurodegenerative disorders, and psychiatric conditions ^1–5^. Redox reactions involve the transfer of electrons from one molecule to another, and these reactions play a fundamental role in energy production, signal transduction, and the maintenance of cellular homeostasis. The majority of NADH is generated from NAD^+^ reduction during glycolysis in the cytosol and the tricarboxylic acid (TCA) cycle in mitochondria. Subsequently, NADH is oxidized to NAD^+^ in the electron transport chain (ETC) ^6^ where the coupling of NADH oxidation and adenosine diphosphate (ADP) phosphorylation leads to substantial adenosine triphosphate (ATP) production. Essential metabolic pathways, including glycolysis, the TCA cycle, ETC, fatty acid oxidation, and ketone body metabolism, heavily depend on redox reactions, with NAD^+^/NADH serving as an important cofactor. Therefore, the redox state of the cellular environment is reflected by NAD^+^ and NADH levels and a disturbance in their homeostasis is likely to impact these metabolic pathways.

Increased neural activity requires high energy supply to sustain neurotransmission. This energy demand is thought to be supported by a larger increase in non-oxidative than oxidative metabolism, as evidenced by a decrease in the oxygen-to-glucose index (OGI) during brief brain stimulation ^7^. Recent findings suggest that astrocytic glycogenolysis plays a compensatory role during these periods of heightened energy demand, sparing glucose for neuronal activity while supporting transient ATP-needs through glycolysis ^8,9^. As NAD^+^ and NADH are critical cofactors involved in brain energy metabolism, their levels are expected to change during sensory stimulation. Controversial results regarding NAD^+^ reduction and NADH oxidation have been reported upon electric stimulation in early studies using optical techniques ^10,11^.

Functional Magnetic Resonance Spectroscopy (fMRS) offers a non-invasive way to study biochemical changes during brain activation *in vivo*. For this, metabolite concentrations are compared between stimulated and baseline states to reveal task-induced changes. ^1^H-fMRS studies in the visual cortex have shown increases in metabolite levels like lactate, and glutamate by about 2–4% during visual stimulation ^12–17, 66^.

In turn, ^31^P MRS is particularly suited to non-invasively assess energy-metabolism in the human brain through the detection of phosphocreatine (PCr), and nucleotides such as ATP, NAD^+^ and NADH ^5,18^. Thus, ^31^P fMRS offers the means to study dynamic responses of NAD^+^ and its reduced form NADH *in vivo*, providing insights into the biochemical basis of brain energy metabolism to support neuronal functioning.

So far, only a limited number of studies have explored *dynamic* NAD^+^ changes in the brain upon sensory stimulation using ^31^P fMRS. Previous studies at 3 Tesla (T) field strength have found no changes in total NAD (tNAD) (with a trend towards a decrease of -5%) ^19^, and a decrease in tNAD ^20^. Notably, spectral resolution at such low magnetic field strengths is insufficient to distinguish NAD^+^ and NADH signals ^20^. Others have used ^31^P MRS at 7T to successfully measure NAD^+^ and NADH *in vivo* ^21^. More recently, Pohmann et al.^62^ employed 9.4T ^31^P MRSI with high spectral resolution and found no significant changes in NAD□ or NADH during visual stimulation. However, dynamic assessments of NAD(H) fluctuations in the human brain remain technically challenging due to both inherently low ^31^P MRS sensitivity, and small effect sizes ^22^. Those challenges can be addressed by studying a large number of participants, and working at the highest sensitivity available.

We therefore investigated NAD=/NADH dynamics in the human occipital lobe during a visual stimulation task using block-design ^31^P fMRS at 7T. Our single-voxel, time-resolved approach allowed block-specific assessment for the first time. Based on previous work, we hypothesized a reduction of NAD□ concentrations during stimulation, accompanied by stable, unchanged ATP concentrations ^20^.

## METHODS AND MATERIAL

### Participants

30 participants (age = 25 ± 4 years, 15 females) were recruited via advertisements at the University campus of the Swiss Federal Technology Institute of Lausanne (EPFL). Inclusion criteria were: age 18–35 years old, with normal or corrected vision and hearing abilities; exclusion criteria were: a history of psychiatric disorders, excessive smoking (>1 pack/day), excessive alcohol consumption (>21 units/week), or other regular drug use, and contraindications to magnetic resonance imaging (MRI) examination.

The study was approved by the Ethics Committee of Canton Vaud, Switzerland (BASEC-2023-00370) and conducted in accordance with the Declaration of Helsinki (latest revision, 2013). All participants provided written informed consent prior to participation.

### MR Measurements

MR experiments were performed on a 7T/68 cm scanner (Siemens Healthineers, Erlangen, Germany) with an in-house-built ^1^H quadrature surface coil (10-cm diameter) and a single-loop ^31^P coil (7-cm diameter) for the coverage of the human occipital lobe. B_0_ field inhomogeneity was optimized in a voxel of interest (VOI) (50×30×40 mm^3^) using first- and second-order shimming (FASTMAP ^23^).

Magnetization Prepared 2 Rapid Acquisition Gradient Echo (MP2RAGE) ^24^ images (TR/TE = 5500/2.3 ms, voxel size = 1 mm^3^ isotropic) were acquired for anatomical localization, to position a 55×20×25 mm^3^ VOI for the MRS measurement in the occipital lobe. Before the fMRS acquisition, a localizer functional MRI (fMRI) scan (3D-EPI, TR/TE = 2820/41.6 ms, volumes = 180, voxel size = 1.9×1.9×2.6 mm^3^) was obtained for accurate MRS voxel placement in the activated occipital area (Figure 1A). Localized ^31^P MR spectra were acquired using a 3D-ISIS sequence (TR/TE = 3000/0.35 ms, voxel-size = 55×20×25 mm^3^, averages/block = 16/6, bandwidth = 6000 Hz, number of points = 2048). The complete measurement details can be found in the “MRSinMRS” table (Supplementary Table 1) ^25,26^.

**Figure 1.**
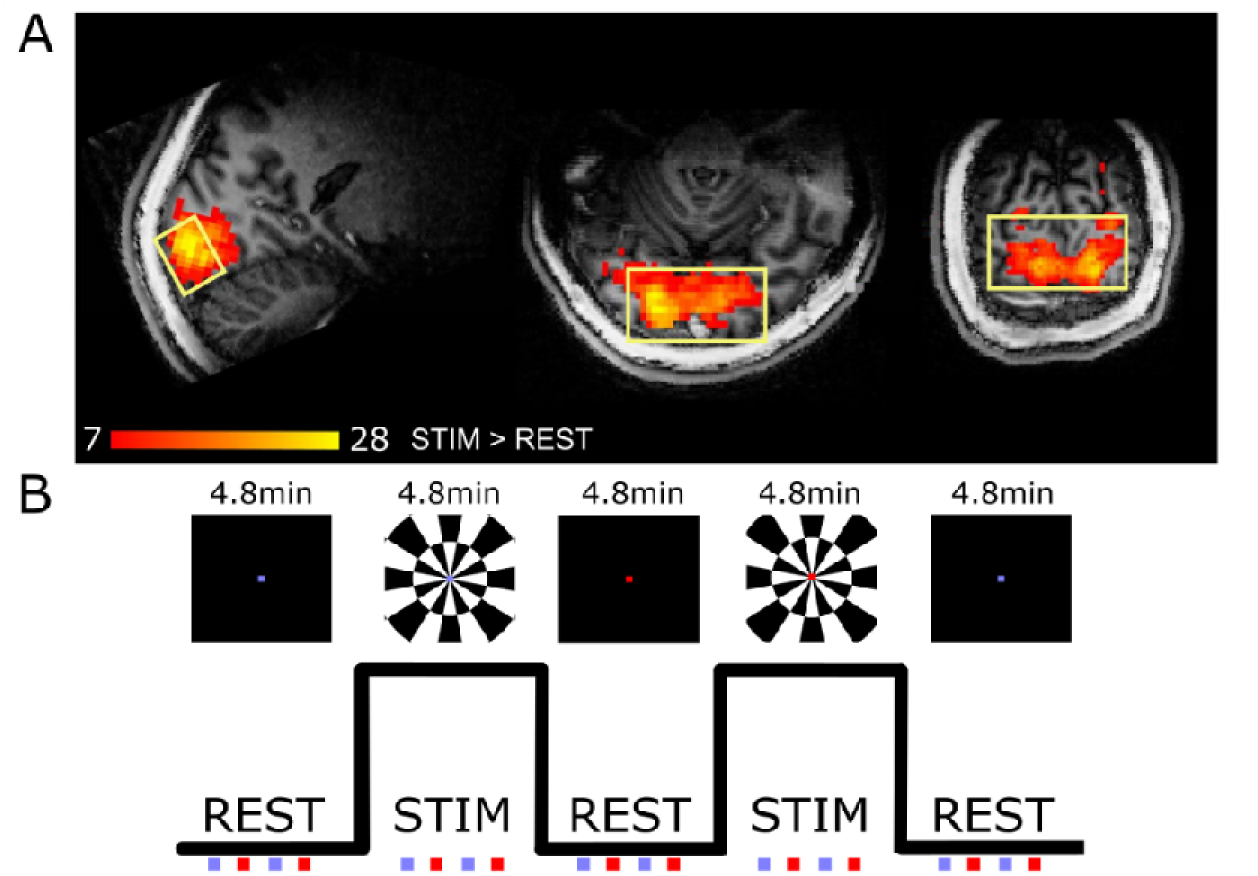
Study design. **A)** A representative fMRI activation-map of the stimulation>rest contrast (red/yellow) and voxel (yellow) overlaid on a representative anatomical image (color bar represents z-scores). **B)** Schematic representation of the stimulation paradigm. As visual stimulation checkerboards were used, in the rest condition a black screen with a fixation dot was used. In both conditions, the fixation dot changed color four times per block. Each block lasted 4.8 min corresponding to 6×16 averages in MRS. The participant was asked to respond to the color change with a button press, as depicted below the paradigm.

The total acquisition time was under 50 min including 15 min for preparation procedures (MRI localizer images, shimming and pulse power calibration), 9 min for the fMRI localizer, 24 min for the dynamic ^31^P MRS acquisition.

### ^31^P fMRS experimental setup

Participants were exposed to an established visual stimulation task (24 min) consisting of alternating blocks of rest (4.8 min) and flashing checkerboard stimuli (4.8 min) ^27^ (Figure 1B), while fMRS spectra were collected from a VOI located in the visual cortex (Figure 1A). Participants viewed the stimuli through a mirror on an LCD screen (viewing distance: ∼160 cm, screen size: 700×395 mm). Visual stimuli were full-field checkerboards, contrast reversing at 8 Hz (100% contrast). In the rest condition, a black screen with a colored fixation dot was used. To ensure attention to the screen, in both conditions, the fixation dot changed color four times per block (total of 20). The participants were asked to respond to the color change with a button press (all button presses summed up /total color changes = response rate). Additionally, the time between button presses was recorded to determine which reactions were on time (response time). Button presses were determined to be invalid if the duration between them was under 70 ms or over 75 ms. Participants were excluded from the analysis when the response rate was lower than 17 out of 20 button presses.

### fMRI localizer setup and analysis

Participants were exposed to a short version of the fMRS experimental setup (4.3 min), with alternating blocks of rest (24s) and flashing checkerboard (24s) while the fMRI scans were conducted. The online fMRI analysis was performed with the Siemens Syngo inline BOLD package, which implements a standard GLM contrast of stimulation versus rest. The resulting activation map was automatically generated on the scanner and used to guide voxel placement for the ^31^P MRS acquisition.

FMRI data were also analyzed subsequently/offline: preprocessing included motion correction (FLIRT), spatial smoothing (5 mm FWHM) and high pass-filtering (100 s) using FSL. FMRI data were entered into the first-level analysis (FSL/FEAT v.6.00; RRID: SCR_002823) ^28^. The model was designed to estimate the effect of visual stimulation on the occipital lobe. To explore whole-volume activity in the main task contrast (stimulation vs. rest), the first-level contrast-of-parameter-estimates (COPE) maps were analyzed using non-parametric permutation testing (5000 permutations), using FSL Randomise. Thresholds for all analyses were initially set at p < 0.05 with family-wise error corrections using threshold-free cluster enhancement ^29^.

### ^31^P MRS preprocessing

Signal to Noise Ratio (SNR) was calculated by LCModel ^30^ dividing the amplitude of PCr by the fitting residual. Linewidth was calculated with the linewidth function of FID-A ^31^, which estimates the full width at half maximum (FWHM) of spectra by calculating the linewidth of PCr in the frequency domain using a Lorentzian fit. Each ^31^P MR spectrum was visually inspected, to ensure that spectra were free of obvious artefacts such as severe baseline distortions, frequency misalignment, or excessive noise. Additionally, data points with a Cramér–Rao lower bound (CRLB) >50% were excluded.

Since the acquisition time was significantly longer than the FID decay time, 5-Hz Lorentzian apodization was applied to the data. All transients were averaged per block after frequency and phase correction, using in-house developed MATLAB (version R2024a, Mathworks, USA) scripts. Signals were phased according to the phosphocreatine (PCr) resonance in the frequency domain.

Additionally, we analyzed the same dataset with the same preprocessing steps plus also applying Marchenko-Pastur principal component analysis (MP-PCA) denoising ^32^ (Supplementary Materials).

### ^31^P MRS quantification

^31^P MR spectra were analyzed using LCModel ^30^ with a basis-set composed of simulated ^31^P spectra of PCr, alpha(α)-, beta(β)-, gamma(γ)-adenosine triphosphate (ATP), intracellular inorganic phosphate (Pi_int_), extracellular inorganic phosphate (Pi_ext_), phosphoethanolamine (PE), phosphocholine (PC), glycerophosphocholine (GPC), glycerophosphoethanolamine (GPE), membrane phospholipid (MP), oxidized Nicotinamide Adenine Dinucleotide (NAD^+^), and its reduced form NADH, and the right resonance at -9.3 ppm of uridine diphosphoglucose (UDPG) with their respective linewidth ^33,34^ (control parameters and basis set composition plot can be found in Supplementary Materials, Supplementary Figure 1). Intracellular (Pi_int_) and extracellular (Pi_ext_) inorganic phosphate resonances were included as separate entries in the LCModel basis set and quantified independently, without coupling or frequency boundary constraints between them. α-ATP, β-ATP and γ-ATP are simulated respectively, as multiplets with J-coupling constant ^69^.

The quantification of NADH in the presence of UDP-glucose (UDPG) has its limitations: resonances at −9.83 ppm and −8.23 ppm are overlapping with NADH and NAD^+^. Only the −9.83 ppm resonance was modeled in this study, as including both doublets at the available SNR led to unstable fits and greater variability. Results did not differ systematically between one- and two-doublet modeling, but the one-doublet strategy yielded more stable estimates (more details in the Supplementary Methods). Accordingly, the reported NADH values should be regarded as NADH+, reflecting partial UDPG contribution.

γ-ATP was assumed to be 2.8 mmol/L in the human occipital lobe and used as an internal concentration reference for all metabolites except NAD^+^, NADH+, tNAD, and UDPG ^35^. To be comparable with literature investigating NAD^+^ and NADH+ concentrations, they were calculated assuming α-ATP of 2.8 mmol/L as an internal concentration reference ^5,36^. Total ATP was calculated as an average of α-ATP, β-ATP and γ-ATP, and referenced to the sum of all metabolites (tP) to investigate the stability of ATP across the experiment. Intracellular pH values were calculated from the chemical shift difference between Pi_int_ and PCr and [Mg^2+^] was calculated from the chemical shift difference between β-ATP and PCr ^37,38^.

Changes in NAD^+^, and total NAD (tNAD = NAD^+^ + NADH+) were statistically tested as the primary endpoints and NADH+, ATP, PCr, Pi_int_, PCr/Pi_int_, pH and [Mg^2+^] were tested as secondary, exploratory endpoint (more information in Supplementary Materials).

## Statistical Analysis

### Power analysis

First, we determined the sample size to assess a hypothesized effect on neurometabolic signaling. A previous study investigating the effects of visual stimulation on glutamate concentration changes using ^1^H-MRS reported an effect size of d = 1.32 (Cohen d) in young participants ^17^. As this study used ^31^P-MRS, we expected a more moderate effect size of d = 0.8. To detect the effect of the task on metabolite concentration changes, with a power of 95% and an α of 0.05, we would need 20 participants.

### Statistical tests

Statistical analyses were conducted using R v.4.3.1 ^39^. Linear mixed-effects models were used to analyze changes in fMRS metabolite concentrations to investigate the main effect of stimulation block (rest-block1 (rest1), stimulation-block1 (stim1), rest-block2 (rest2), stimulation-block2 (stim2), rest-block3 (rest3); here called “BLOCK”), with linewidth as a covariate using the lme4 package ^40^. Multiple comparison correction over all primary metabolite change tests was performed using Sidak’s correction: *α*^*^ = 1 - (1 - *α*)^1/*m*^, with α = 0.05 and m = 3 (number of tests for the primary hypothesis), which resulted in an α* = 0.02. Additionally, Tukey corrected t-tests were used as post-hoc tests to test the effect of the respective single blocks.

Furthermore, we conducted exploratory analyses where the blocks were split into two halves, allowing us to observe more granular metabolic changes within each block (see Supplementary Methods). Data is shown as mean ± SD, unless otherwise stated.

## RESULTS

### Participants

Four participants were excluded based on the button presses (low response rate) during the experiment, indicating that no attention was paid to the task, and one because of low overall SNR (<12), resulting in twenty-five datasets (age = 24 ± 4 years, 10 female) that were included in the analysis. The average response rate over all participants was 100%, with a mean response time of 71.4 ± 6.0 ms.

### FMRI localizer analysis

fMRI analysis showed a significant BOLD effect of stimulation vs. fixation in the visual cortex located in the occipital lobe (Figure 1). When overlaying the spectroscopy voxel with the activation map, a high overlap was found, indicating that the MRS measurement voxel was placed in a brain region showing high BOLD signal in response to the stimulation, as intended.

## fMRS data

### Quality Measures

^31^P spectral linewidth (LW) did not show any significant changes over the course of the experiment (F(4,88) = 1.69, p = 0.16), with a mean of 12.4 ± 0.4 Hz, indicating good quality and stability. The water linewidth in the VOI was 16.8 ± 1.3 Hz. The mean SNR of ^31^P MR spectra over all blocks was 16 ± 1, no transients of the included datasets were discarded based on visual quality control. The averaged spectra per block showed visible changes per condition when zoomed into the spectral range around NAD (Figure 2A).

**Figure 2.**
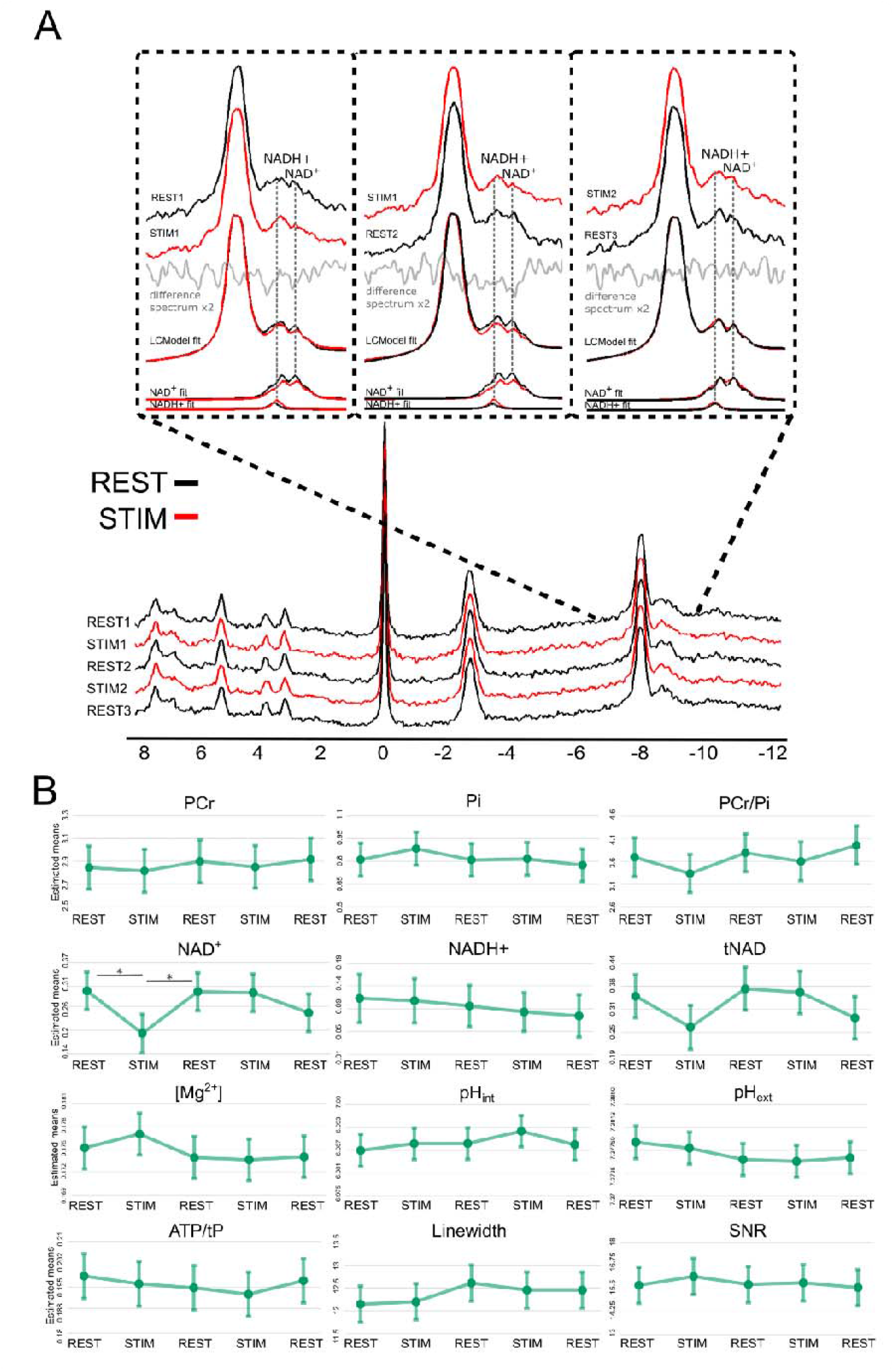
Results. **A)** Quality measures: The sum of all spectra of all included participants during the rest (black) and activation (red) blocks were averaged together and plotted above each other, with a highlight on the NAD^+^ area around -9ppm, including a difference-spectrum of rest-activation (gray), and the NAD^+^ and NADH+ fits by LCModel. **B)** Individual concentration changes (estimated means are presented alongside their 95% confidence intervals (CI)) per block (5-block analysis; rest1, stim1, rest2, stim2, rest3) for PCr, Pi, PCr/Pi, NAD^+^, NADH+, tNAD, [Mg2+], pH_int_, pH_ext_, and ATP/tP, and Linewidth, and Signal-to-Noise-Ratio (SNR). * = p<0.02

CRLBs per fitted metabolite are shown in Supplementary Table 3. All concentrations were in the range of those previously reported (Figure 2 and Table 1) ^19,34^. NADH+ concentration and PH_ext_ could only be obtained for ∼50% of the time points (more information in Supplementary Results). This is due to their low concentrations and high overlap with the NAD^+^ and PH_int_ peaks, respectively. A zoomed in fit-quality plot for Pi can be found in Supplementary Figure 6.

**Table 1.** Statistical analysis. The results of the main analysis and post-hoc tests conducted in our study are shown. The analysis was performed using linear mixed effects models to estimate the means and confidence intervals for each parameter. Post-hoc comparisons were conducted using Tukey’s HSD tests. The parameters investigated include linewidth and various metabolites and physiological parameters. The estimated means are presented alongside their 95% confidence intervals (CI) per block and associated statistical significance values (p-values). Concentrations of NAD^+^, NADH+, tNAD, UDPG are given in mM, referenced to an assumed =-ATP concentration of 2.8 mM, PCr, Piint, are given in mM, referenced to an assumed γ-ATP concentration of 2.8 mM in the human occipital lobe. [Mg^2+^] is reported in mM. ATP was calculated as an average of α-ATP, β-ATP and γ-ATP, and referenced to the sum of all metabolites (tP). Table1. Statistical analysis. The results of the main analysis and post-hoc tests conducted in our study are shown. The analysis was performed using linear mixed effects models to estimate the means and confidence intervals for each parameter. Post-hoc comparisons were conducted using Tukey’s HSD tests. The parameters investigated include linewidth and various metabolites and physiological parameters. The estimated means are presented alongside their 95% confidence intervals (CI) per block and associated statistical significance values (p-values). Concentrations of NAD^+^, NADH+, tNAD, UDPG are given in mM, referenced to an assumed =-ATP concentration of 2.8 mM, PCr, Piint, are given in mM, referenced to an assumed γ-ATP concentration of 2.8 mM in the human occipital lobe. [Mg^2+^] is reported in mM. ATP was calculated as an average of α-ATP, β-ATP and γ-ATP, and referenced to the sum of all metabolites (tP).

| Metabolite | Estimated mean parameters [95% CIs] |  |  |  |  | Statistics (block+linewidth) | Post-Hoc test |
| --- | --- | --- | --- | --- | --- | --- | --- |
|  | Rest 1 | Stim 1 | Rest 2 | Stim 2 | Rest 3 |  |  |
| Linewidth (Hz) | 12.77 [11.75 12.53] | 12.19 [11.80 12.59] | 12.61 [12.21 13.00] | 12.45 [12.06 12.85] | 12.45 [12.06 12.85] | F(4,88) = 1.69, p = 0.16 | - |
| PCr (mM) | 2.84 [2.65 3.03] | 2.82 [2.63 3.00] | 2.90 [2.71 3.09] | 2.85 [2.66 3.04] | 2.92 [2.73 3.10] | F(4,88) = 0.24, p = 0.92 | - |
| Pi<sub>int</sub> (mM) | 0.81 [0.70 0.92] | 0.88 [0.78 0.99] | 0.81 [0.70 0.92] | 0.82 [0.71 0.93] | 0.77 [0.67 0.88] | F(4,88) = 0.57, p = 0.69 | - |
| PCr/Pi<sub>int</sub> (-) | 3.71 [3.27 4.13] | 3.35 [2.92 3.77] | 3.79 [3.37 4.21] | 3.59 [3.17 4.02] | 3.95 [3.53 4.38] | F(4,86) = 1.12, p = 0.35 | - |
| NAD<sup>+</sup> (mM) | 0.29 [0.25 0.35] | 0.19 [0.14 0.24] | 0.29 [0.24 0.34] | 0.29 [0.25 0.34] | 0.24 [0.19 0.29] | F(4,83) = 4.39, p<0.01 | REST1>STIM1 t(83) = 3.38, p<0.01;<br>STIM1<REST2 t(83) = -3.29, p = 0.01 |
| NADH<sup>+</sup> (mM) | 0.12 [0.06 0.17] | 0.11 [0.06 0.16] | 0.10 [0.06 0.14] | 0.09 [0.05 0.13] | 0.04 [0.05 0.12] | F(4,60) = 0.45, p = 0.77 | - |
| tNAD (mM) | 0.35 [0.29 0.41] | 0.27 [0.21 0.33] | 0.37 [0.31 0.43] | 0.36 [0.30 0.42] | 0.29 [0.23 0.35] | F(4,83) = 2.54, p = 0.05 | - |
| pH<sub>int</sub> | 6.987 [6.983 6.992] | 6.989 [6.985 6.994] | 6.989 [6.985 6.994] | 6.992 [6.988 6.997] | 6.989 [6.985 6.993] | F(4,88) = 0.76, p = 0.56 | - |
| pH<sub>ext</sub> | 7.378 [7.376 7.381] | 7.378 [7.375 7.380] | 7.377 [7.373 7.379] | 7.376 [7.373 7.378] | 7.376 [7.374 7.379] | F(4,88) = 0.99, p = 0.42 | - |
| [Mg<sup>2+</sup>] (mM) | 0.175 [0.172 0.178] | 0.177 [0.174 0.179] | 0.174 [0.171 0.176] | 0.174 [0.170 0.176] | 0.174 [0.171 0.177] | F(4,88) = 1.27, p = 0.29 | - |
| ATP/tP (-) | 0.198 [0.191 0.206] | 0.196 [0.189 0.204] | 0.195 [0.187 0.202] | 0.193 [0.185 0.199] | 0.197 [0.189 0.204] | F(4,88) = 0.59, p = 0.67 | - |
| UDPG (mM) | 0.32 [0.24 0.39] | 0.29 [0.21 0.38] | 0.29 [0.21 0.38] | 0.33 [0.23 0.42] | 0.35 [0.26 0.44] | F(4,39) = 0.28, p = 0.89 | - |

### Metabolite changes

We have reliably determined concentrations of PCr, Pi_int_, ATPs, NAD^+^ and tNAD from ^31^P spectra acquired in 4.8 minutes (Table 1). In addition, the measured spectra allowed us to estimate intra- and extracellular pH and intracellular magnesium levels (Table 1). NAD^+^ showed significant changes in response to the visual stimulation, with a significant main effect of block (p<0.01). The effect was driven by a significant concentration decrease during the first stimulation and subsequent symmetric recovery (log-change ±0.42), consistent with a transient 35% dip during the initial stimulation block. NADH+ concentrations remained unaltered, therefore total NAD (tNAD) followed a similar oscillation as observed in NAD^+^ (log-change ±0.260, corresponding to 23% decrease during the first stimulation block). tNAD showed a strong effect of block, which did not survive the multiple comparison correction (p = 0.05). PCr, Pi_int_, and PCr/Pi_int_, NADH+, ATPs, pH and [Mg^2+^] did not show any significant changes over the course of the experiment (all p>0.05). All results of the statistical tests, estimated means, and 95% confidence intervals can be found in Table 1.

Exploratory analysis by splitting the blocks into halves, resulting in 2.4min of acquisition, are detailed in Supplementary Figure 7 and the accompanying table (Supplementary Table 4). Although not statistically significant, likely due to limited power, there appears to be a trend toward a reduction in NAD^+^ and tNAD levels, particularly in the second half of both stimulation blocks.

Application of MP-PCA to the spectral data yielded the same results (Supplementary Figure 9). All statistical tests, estimated means, and 95% confidence intervals can be found in the Supplementary Results.

## DISCUSSION

We have demonstrated the feasibility of measuring time-resolved, block-specific NAD^+^ dynamics in the human occipital lobe during visual stimulation for the first time, using ultra-high field ^31^P fMRS. Our findings revealed a stimulation-induced reduction of 35% in NAD^+^ levels, without significant changes in NADH+, therefore resulting in a trend of 23% lowering of total NAD (tNAD) levels during the first visual stimulation block. While initially it appeared that there was no further reduction in NAD^+^ during the second stimulation block, exploratory analyses hint at a trend also towards a continued reduction in the second halves of the stimulation blocks (Supplementary Table 4, Supplementary Figure 7).

### NAD^+^ Reduction During Stimulation

NAD^+^ and NADH are central to energy metabolism. NAD^+^ acts as a cofactor for glyceraldehyde-3-phosphate dehydrogenase (GAPDH) during glycolysis generating NADH. After glycolysis, cytosolic pyruvate may be converted to lactate by lactate dehydrogenase (LDH), simultaneously oxidizing NADH back to NAD^+^. Alternatively, cytosolic NADH can be oxidized to NAD^+^ through the malate-aspartate shuttle (MAS) ^41^, which transfers reducing equivalents from cytosol to mitochondria. These mechanisms ensure the maintenance of cytosolic NAD^+^ homeostasis and an efficient glycolysis ^42^ (Figure 3).

**Figure 3.**
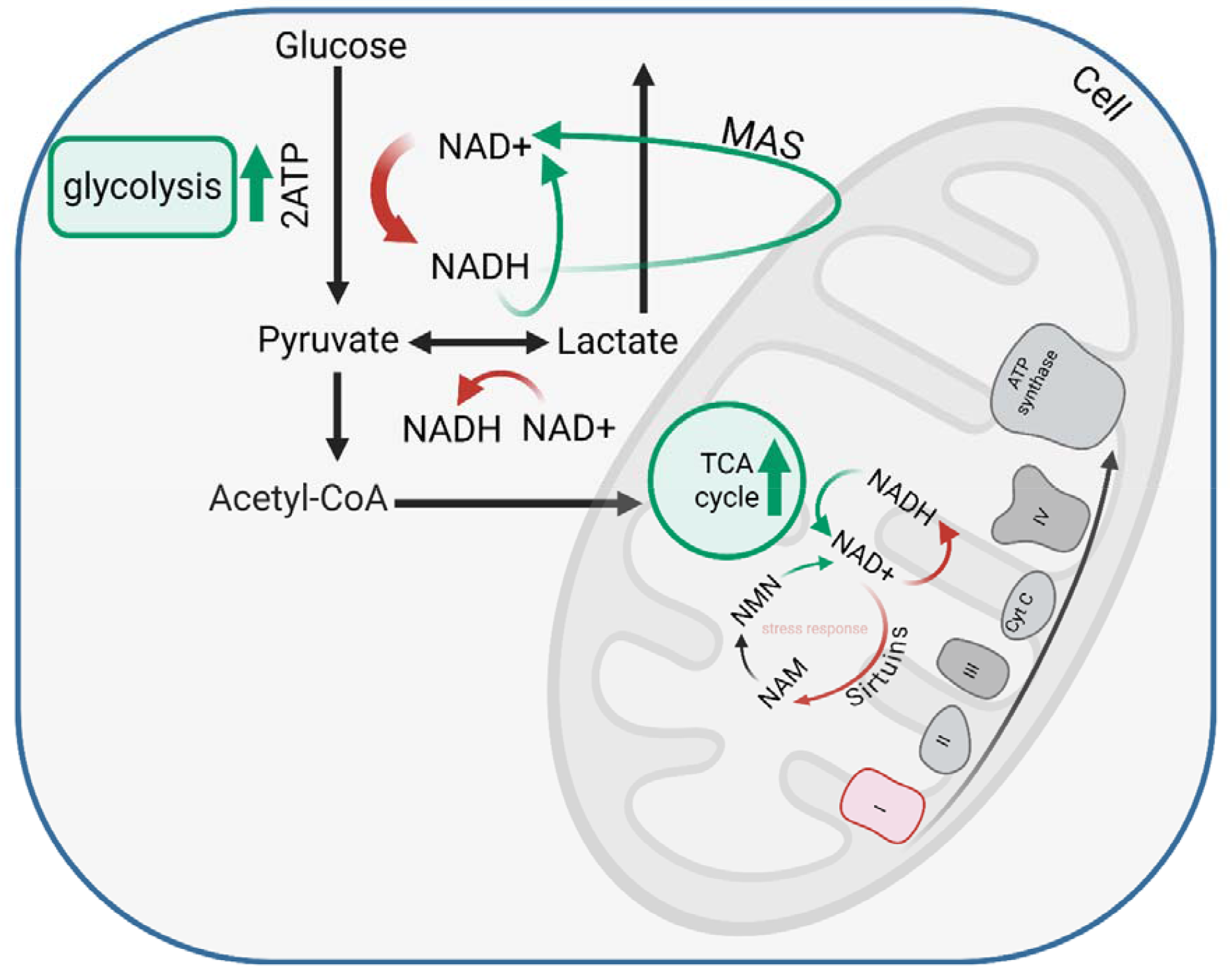
Overview of the NAD-related pathways during brain stimuli. The stimulation-induced NAD^+^ response illustrates a complex metabolic interplay during cortical activation. During the initial visual stimulation block, as depicted in the figure, increased glycolysis accelerates the reduction of NAD^+^, likely exceeding the capacity of oxidative phosphorylation and the malate-aspartate shuttle (MAS) to regenerate NAD^+^. This rapid consumption of NAD^+^ is further compounded by sirtuin activation, which utilizes NAD^+^ to counteract metabolic stress. However, contrary to initial observations suggesting a stabilization of NAD^+^ levels, exploratory analysis indicates a continuing trend towards NAD^+^ reduction in the second halves of the stimulation blocks, suggesting that the adaptive mechanisms—such as enhanced mitochondrial NAD^+^ regeneration, AMPK-mediated metabolic adjustments, and lactate-driven inhibition of cAMP signaling—might not fully restore NAD^+^ homeostasis within the duration of the blocks. This implies that the metabolic adjustments during the later phases of stimulation are more nuanced and may require extended periods to stabilize.

For the first time, we observed dynamic changes in NAD^+^ during visual stimulation, noting a pronounced reduction in NAD^+^ particularly in the initial stimulation block. This observation suggests that the metabolic demand during high activity phases stimulates glycolysis at a rate that outpaces NAD^+^ replenishment via LDH and/or MAS activity. In an earlier study at a lower magnetic field, Barreto et al. ^20^ reported a 2% decrease in total NAD (tNAD) levels during a similar 5-minute stimulation period, aligning with our findings. Our results differ from those of Pohmann *et al*.^62^, who, using 9.4T ^31^P MRSI with high spectral resolution, reported no significant changes in NAD□ or NADH during visual stimulation. A key difference is that their analysis was based on spectra averaged across repeated rest–stimulation cycles, whereas our single-voxel approach revealed a block-specific decrease in NAD□ during the first but not the second stimulation period. This suggests that methodological differences in temporal resolution and averaging strategy may strongly influence detectability of NAD□ dynamics, highlighting the advantage of measuring a time course to identify time-dependent responses.

Furthermore, others have observed a 3-4% decrease in NADH levels in hippocampal slices, followed by an approximate 4% increase that peaked at 1 minute and was sustained for several minutes during prolonged stimulation (1-4 minutes) ^43^. The sensitivity of ^31^P MRS may not detect such minor fluctuations in NADH given its low level, therefore, the observed reduction in tNAD is primarily attributed to NAD^+^ depletion. This suggests that the transient reduction of NAD^+^ by GAPDH temporarily exceeds its recycling via the conversion of pyruvate to lactate.

In fact, employing an NADH biosensor, Díaz-García et al. demonstrated that transient NADH production from glycolysis exceeds the rate of NADH shuttling to mitochondria ^44^, indicating that increased glycolysis relative to MAS activity leads to a net reduction in NAD^+^. On the other hand, previous ^1^H fMRS studies have consistently reported increased lactate concentration during visual stimuli, suggesting the heightened LDH activity to recycle NAD^+^ from NADH, thereby supporting glycolysis. The concurrent increase in glutamate, and decrease in aspartate, during sensory stimulation, another replicated finding in ^1^H fMRS studies^15,17,45^, may result from enhanced MAS activity, involving the rate-limiting Glu-Asp antiporter ^45^, or from an increase in Cerebral Metabolic Rate of Oxygen (ΔCMR_O2_) that is below the Cerebral Metabolic Rate of Glucose (ΔCMR_Glc_) ^46^. Notably, in the study by Mangia et al.^45^, the second stimulation block elicited smaller, though still detectable, increases in both lactate and glutamate, suggesting that some degree of attenuation or adaptation across repeated blocks can occur. Furthermore, compartmental shifts of glutamate between pools with differing T□ relaxation times and thus differing MRS visibility have been proposed as a possible explanation, particularly in event-related paradigms at lower fields with long-TE acquisitions^60,61^. In addition to GAPDH-dependent NAD^+^ reduction, the TCA cycle enzymes α-ketoglutarate dehydrogenase, malate dehydrogenase and isocitrate dehydrogenase (IDH) also reduce NAD^+^ to NADH, which provides electrons for proton pumping at the ETC and subsequent oxidative phosphorylation.

Beyond its role as a metabolic redox cofactor, NAD^+^ also serves as an essential substrate for enzymes such as Poly(ADP-ribose) polymerases and sirtuins, which activation contributes to NAD^+^ depletion ^47^. In particular, sirtuins are NAD^+^-dependent enzymes that regulate transcriptional responses to metabolic stress, including the upregulation of antioxidant defenses. Given that strenuous and prolonged stimulation-induced cortical activity is metabolically demanding for neurons and astroglia ^48^, it is plausible that sirtuins are activated to mitigate oxidative stress, consuming NAD^+^ in the process. However, in most tissues, transcript levels of the key glycolytic enzyme GAPDH are several orders of magnitude higher than those of NAD^+^-consuming enzymes, suggesting that glycolysis-driven NAD^+^ reduction might be the predominant NAD^+^ usage.

The mitochondrial NAD^+^ pool can be replenished through direct import from the cytosol or synthesized locally from imported nicotinamide mononucleotide (NMN). Complex I of the ETC is the primary site for oxidizing NADH back to NAD^+^ using oxygen (O_2_). Brain activation leads to increased cerebral blood flow (CBF), CMR_glc_, and CMR_O2_, resulting in higher blood oxygenation ^9,49^. However, a mismatch exists between CMR_glc_ and CMR_O2_, shifting the OGI from rest to activation ^7^. Both blood flow and glucose utilization increase in parallel during neuronal activation, exceeding oxygen utilization. This mismatch in glucose and oxygen use during activation may temporarily slow NAD^+^ regeneration via ETC activity compared to non-oxidative NAD^+^ recycling, leading to transient overall NAD^+^ depletion.

To our surprise, a second period of intense visual stimulation did not result in NAD^+^ reduction, suggesting a local metabolic adaptation. Two pathways appear to be well suited to generate NAD^+^ in such conditions of increased cortical activity. Firstly, NAD^+^ is consumed by sirtuins with formation of nicotinamide that can be used in a salvage pathway to produce the NAD^+^ precursor nicotinamide mononucleotide (NMN). It is known that neuronal activation is accompanied by stimulation of the AMPK pathway ^50^, and AMPK activation can stimulate both sirtuins ^51^ and the NAD^+^ salvage pathway ^52^. Since AMPK-dependent regulation of the salvage pathway requires not only protein phosphorylation but also upregulation of gene expression ^52^, it is likely to be more pronounced in the second but not the first stimulation period of our study. Notably, mitochondrial NAD^+^/NADH redox can be maintained despite significantly enlarged NAD pool upon increased levels of NMN ^53^.

Secondly, NAD^+^ is produced *de novo* from tryptophan via the kynurenine pathway, which also generates a number of neuroactive metabolites ^54^. It has been reported that pyruvate stimulates the release of kynurenic acid from astrocytes ^55^, suggesting that glycolytic stimulation could result in stimulation of the kynurenine pathway and possibly of NAD^+^ synthesis. Kynurenine metabolism is acknowledged to take place in glial cells rather than neurons ^56^. However, to our knowledge it remains unknown whether astrocytic activation during increased cortical stimulation directly induces NAD^+^ synthesis *de novo*.

While these hypotheses are biologically plausible and supported by literature, our data do not directly assess these mechanisms and thus they remain speculative.

### pH Remains Unchanged

The results of pH changes during stimulation are inconsistent with prior studies using ^31^P-MRS. The absence of significant pH changes during stimulation was reported at 3T^20^ and 7T^19^. While our data did not reveal significant pH changes during stimulation, it should be noted that small pH_int_ increases have been reported previously, including a study by Zhu *et al*.^63^ that also found a concomitant decrease in [Mg^2^=]_i_, as well as earlier reports of stimulation-induced alkalinization^67,68^. In line with these findings, Pohmann *et al*.^62^ and Hendriks *et al*.^64^ also reported subtle shifts in Pi resonances during visual stimulation, suggestive of pH modulation. In our dataset, pH_int_ could be quantified reliably and remained stable, whereas pH_ext_ could only be derived in <50% of time points due to the low SNR of Pi_ext_ (see Supplementary Methods and Results), limiting the reliability of detecting stimulation-induced pH_ext_ changes. The stability of pH in our study suggests that any increase in H^+^ and lactate production is effectively buffered by intracellular mechanisms or exported, preventing detectable shifts in pH.

One possible explanation for the observed pH stability is the interaction between glycolysis and astrocytic glycogen metabolism. Studies suggest that during neuronal activation, astrocytes preferentially utilize glycogen-derived glucose rather than extracellular glucose ^8^. Moreover, the activation of AMPK during the first stimulation block could ensure rapid mitochondrial adaptation, preventing sustained lactate accumulation that could otherwise acidify the intracellular environment ^57^.

Additionally, small, non-significant dips in PCr/Pi_int_ were observed during both stimulation blocks. These may reflect transient utilization of PCr to buffer ATP demands. This interpretation is supported by prior findings of decreased PCr/Cr during stimulation^48^. In addition, this observation may mainly result from the tendency of an increase in Pi and a reduction in PCr, which were also observed in the ^31^P fMRS study of Zhu et al.^63^.

### Technical Aspects

The technical aspects of the study played a significant role in ensuring the quality and reliability of the obtained spectra. With a temporal resolution of 4.8 min, a mean spectral SNR of around 16 for each block was consistently maintained during the functional task, suggesting a high stability of the spectra quality over time. This was achieved through FASTMAP shimming^23^ to minimize local field inhomogeneities and spectral registration to minimize frequency and phase variations. Additionally, the use of localized 3D ISIS minimized partial volume effects from non-activated regions, ensuring that the observation of metabolic changes was specific to the stimulated area. One of the key strengths of this study is its implementation at ultra-high field (7T), including a large, and therefore sufficient sample size of 25 participants. High sensitivity, spectral resolution, and spectral quality enable the reporting of the NAD^+^ time course during the visual stimulation paradigm in a block-wise manner. The visual assessment of changes in the NAD^+^ peak, on the right side of the α-ATP resonance peak (magnified part of Figure 2A), confirmed dynamic changes in NAD^+^ levels in response to increased neuronal activity (Figure 2A). Notably, the absence of a BOLD effect on the PCr linewidth and peak height (Figure 2B; p = 0.16, Supplementary Figure 5) suggested minimal vascular contributions to the observed metabolic changes, highlighting the specificity of the findings to metabolic activity.

A limitation concerns the quantification of NADH in the presence of UDP-glucose (UDPG), which has doublets at −9.83 ppm and −8.23 ppm overlapping with NADH and NAD□ ^65^. Only the −9.83 ppm doublet was modeled, as including both doublets at the available SNR led to unstable fits and greater variability. Results did not differ systematically between one- and two-doublet modeling, but the one-doublet strategy yielded more stable estimates (more details in the Supplementary Results). Accordingly, the reported NADH values should be regarded as NADH+, reflecting partial UDPG contribution. In contrast, NAD□ shows a distinct doublet at higher concentration and is more reliably quantified. The absence of block-wise changes in UDPG (Supplementary Figures 3–4) further supports that the observed NAD□ reductions are unlikely to be due to UDPG. Future studies at higher field strength are needed to improve SNR and enable robust separation of NADH and UDPG. The absence of a measurable BOLD effect in our dataset is consistent with previous reports at 9.4 T^63^ but contrasts with Hendriks et al.^64^, who observed a ∼2–3% increase in PCr signal during visual stimulation after averaging across participants. In our data, PCr linewidth remained unchanged (Figure 2B) and no systematic change in PCr peak height was detected (Supplementary Fig. 5). One explanation is that the relatively large voxel sizes used in our experiment, while necessary to ensure sufficient SNR for NAD(H) quantification, encompassed both activated and non-activated tissue (Figure 1A, Supplementary Figure 2), potentially obscuring subtle BOLD-related effects. This is in line with the argument by Pohmann *et al*.^63^ that voxel heterogeneity can dilute small activation-induced spectral changes. Additionally, the expected linewidth change for a ^31^P BOLD effect at 7 T is very small (<0.3 Hz when scaling from ^1^H BOLD linewidth changes), which may be below the sensitivity of our current setup.

Despite the SNR benefit of ultra-high fields, the sensitivity of current techniques for measuring NADH dynamics remains a considerable challenge due to the spectral overlaps with NAD^+^ and α-ATP and the low concentration of the NADH signal. In this study, concentrations for NADH+ could only be extracted in 50% of the timepoints. Similarly, reliable quantification of extracellular pH (pH_ext_) is hampered by the poor detectability of Pi_ext_; less than 50% of the Pi_ext_ fits passed the CRLB < 50% criterion (see Supplementary Table 3). The difficulty in capturing the changes in NADH levels is the current limitation of ^31^P functional MRS, which might be enhanced by advanced denoising methods in the future ^58,59^. We here investigated the MP-PCA denoising method, and obtained the same results with MP-PCA as without, indicating the validity of such methods when applied to ^31^P fMRS data; however, it remains insufficient for detecting NADH(+) changes.

### Clinical Implications

The decline in NAD^+^ availability and abnormal NAD^+^/NADH redox states have also been linked to age-related metabolic diseases and neurodegenerative disorders. The application of high magnetic fields and ^31^P-MRS has enabled direct measurements of these metabolites in the human brain ^3,5,36^. It is essential to clarify that our findings of reduced NAD^+^ during visual stimulation do not directly correspond to static measurements of NAD^+^/NADH ratio in specific patient groups. Instead, dynamic measurements in patient groups could provide nuanced insights into the temporal aspects of NAD^+^ metabolism and its potential implications for cognitive and psychiatric conditions. For instance, investigating how NAD^+^ responds to specific cognitive challenges or stressors over time may offer a more comprehensive understanding of the role of NAD^+^ in the context of cognitive function and psychiatric disorders.

## CONCLUSION

By employing ultra-high field 7T ^31^P fMRS, we demonstrated the feasibility of measuring dynamic NAD^+^ changes in the occipital lobe. Our findings support the sensitivity of fMRS to detect stimulus-induced dynamics in cerebral NAD□ metabolism, providing insights into the interplay between glycolysis and oxidative phosphorylation during neural activation. Our results demonstrate the feasibility of tracking NAD□ dynamics *in vivo* and offer new opportunities for probing energy metabolism in the human brain. While initial observations suggested a subsequent stabilization of NAD^+^ levels, exploratory analyses hint at a trend towards further reductions in the second halves of both stimulation blocks. Further studies with higher sensitivity, temporal resolution and statistical power are needed to confirm and extend these findings.

## ACKNOWLEDGEMENTS

We thank all the participants for contributing to this study, and Silvia Mangia for several fruitful, in-depth discussions about the results.

## AUTHOR CONTRIBUTION STATEMENT

AK, and LX developed the idea and experimental setup, AK, and FAV collected and analyzed the data, MW, and DW developed hardware including the head coil, AK, and LX wrote the manuscript with the inputs from all co-authors (FAV, JD, IJ, YL, XY, MW, DW). JD provided insights for the interpretation of the results and revised the manuscript. IJ provided insights and methods for the MP-PCA approach.

## ABBREVIATIONS

AMPK: AMP-activated Protein Kinase
ATP: Adenosine Triphosphate
CBF: Cerebral Blood Flow
CMR_glc_: Cerebral Metabolic Rate of Glucose
CMR_O2_: Cerebral Metabolic Rate of Oxygen
COPE: Contrast of Parameter Estimates
CRLB: Cramer-Rao Lower Bound
cAMP: Cyclic Adenosine Monophosphate
ETC: Electron Transport Chain
FID: Free Induction Decay
FID-A: FID Appliance
FLIRT: FMRIB’s Linear Image Registration Tool
FSL: FMRIB Software Library
FWHM: Full Width at Half Maximum
GPC: Glycerophosphocholine
GPE: Glycerophosphoethanolamine
GSG: Glucose Sparing by Glycogenolysis
LCModel: Linear Combination of Model spectra
MAS: Malate-Aspartate Shuttle
MP-PCA: Marchenko-Pastur Principal Component Analysis
MRI: Magnetic Resonance Imaging
NAD^+^: Nicotinamide adenine dinucleotide (oxidized form)
NADH: Nicotinamide adenine dinucleotide (reduced form)
NADH+: Nicotinamide adenine dinucleotide (reduced form) with contributions of UDPG
OGI: Oxygen-Glucose Index
OxPhos: Oxidative Phosphorylation
PC: Phosphocholine
PCr: Phosphocreatine
PE: Phosphoethanolamine
Pi_int_: Intracellular Inorganic Phosphate
P_extt_: Extracellular Inorganic Phosphate
SNR: Signal to Noise Ratio
TCA: Tricarboxylic Acid Cycle
tNAD: Total NAD (NAD^+^ plus NADH+)
UPDG: Uridine Diphosphoglucose
fMRS: Functional Magnetic Resonance Spectroscopy
^31^P: Phosphorus-31

