## Supplementary for "Ultra-High Field ^31^P functional Magnetic Resonance Spectroscopy Reveals NAD^+^ Dynamics in Brain Energy Metabolism during Visual Stimulation"

### Supplementary Materials:

Calculation of physiological parameters including pH and [Mg2+]

$$pH={pK}_{a}+ log \frac{\delta_{Pi} - \delta_{a}}{\delta_{b} - \delta_{Pi}}$$

where δPi is the chemical shift difference between PCr and Pi; pKa = 6.73, δ_a_ = 3.275, δ_b_ = 5.385. [Mg^2+^] was calculated from the chemical shift difference between β-ATP and PCr ^1^:

$${pMg}^{2+}= 4.24 - log \left[ \frac{{(\delta_{\beta ATP}+ 18.58)}^{0.42}}{({-15.74 - \delta_{\beta}ATP)}^{0.84}} \right]$$

$$[{Mg}^{2+}]= {{log}_{10}}^{{-pMg}^{2+}}$$

where δ_βATP_ is the difference in chemical shift between β-ATP and PCr.

**Supplementary Table 1**: MRSinMRS checklist

| 1. Hardware |  |
| --- | --- |
| a. Field strength [T] | 6.98 T |
| b. Manufacturer | Siemens Healthineers, Erlangen, Germany |
| c. Model (software version if available) | Magnetom 7T (VB17) |
| d. RF coils: nuclei (transmit/receive), number of channels, type, body part | in-house-built 1H quadrature surface coil (10 cm-diameter) and a single-loop ^31^P coil (7 cm-diameter) for the coverage of the human occipital lobe |
| e. Additional hardware |  |
| 2. Acquisition |  |
| a. Pulse sequence | 3D ISIS localization sequence |
| b. Volume of interest and VOI locations | Single voxel placed in occipital lobe, across hemispheres |
| c. Nominal VOI size [cm^3^, mm^3^] | 55x20x25mm^3^ |
| d. Repetition Time (TR), Echo Time (TE) [ms, s] | TR = 3 s, TE = 0.35 ms |
| e. Total number of excitations or acquisitions per spectrum (NA)  In time series for kinetic studies  i. Number of averaged spectra) per time-point (NA)  ii. Averaging method (e.g. block-wise or moving average)  Total number of spectra (acquired / in time-series) | *16 averages per spectrum (NA = 16). Total number of time points in the time series was 30, at NA = 16, with 6 time points per block, and 5 blocks in total.*  *ii. Averaging method = per block* |
| f. Additional sequence parameters (spectral width in Hz, number of spectral points, frequency offsets)  i. If STEAM:, Mixing Time (TM)  ii. If MRSI: 2D or 3D, FOV in all directions, matrix size, acceleration factors, sampling method | 6000Hz, 2048 complex points |
| g. Water suppression method | n.a. |
| h. Shimming method, reference peak, and thresholds for “acceptance of shim” chosen | *1st and 2nd order shimming using FAST(EST)MAP ^3^, line-width of PCr peak was evaluated post-hoc* |
| i. Triggering or motion correction method  (respiratory, peripheral, cardiac triggering, incl. device used and delays) |  |
| 3. Data analysis methods and outputs |  |
| a. Analysis software | ^31^P MR spectra were visually inspected, and transients with visually defined low signal-to-noise ratio (SNR) would be discarded. Consequently, the remaining transients were summed after frequency and phase correction, using in-house developed MATLAB scripts. Signals were phased according to the PCr resonance in the frequency domain^13^ . 5Hz Lorentzian apodization was applied to the data.  ^31^P MR spectra were analyzed using LCModel ^2^ with a basis-set composed of simulated ^31^P spectra of phosphocreatine (PCr), alpha Adenosine Triphosphate (α-ATP), beta Adenosine Triphosphate (β-ATP), gamma Adenosine Triphosphate (γ-ATP), internal inorganic phosphate (Piint), external inorganic phosphate (Piext), Phosphoethanolamine (PE), phosphocholine (PC), Glycerophosphocholine (GPC), Glycerophosphoethanolamine (GPE), Monophosphate (MP), Nicotinamide Adenine Dinucleotide (NAD^+^), and its reduced form NADH, and uridine diphosphoglucose (UDPG) with their respective linewidth ^11,12^ . |
| b. Processing steps deviating from quoted reference or product analysis software (vendor, version) |  |
| c. Output measure  (e.g. absolute concentration, institutional units, ratio) Processing steps deviating from quoted reference or product |  |
| d. Quantification references and assumptions, fitting model assumptions | *The quantification of NADH in the presence of UDP-glucose (UDPG) has its limitations: resonances at −9.83 ppm and −8.23 ppm are overlapping with NADH and NAD⁺. Only the −9.83 ppm resonance was modeled in this study, as including both doublets at the available SNR led to unstable fits and greater variability. Results did not differ systematically between one- and two-doublet modeling, but the one-doublet strategy yielded more stable estimates (more details in the Supplementary Methods). Accordingly, the reported NADH values should be regarded as NADH+, reflecting partial UDPG contribution.*  *γ-ATP was assumed to be 2.8 mM in the human occipital lobe and used as an internal concentration reference for all metabolites except NAD^+^, NADH+, and tNAD. To be comparable with literature investigating NAD^+^ and NADH concentrations, they were calculated assuming α-ATP of 2.8 mM as an internal concentration reference ^6,7^. ATP was calculated as an average of α-ATP, β-ATP and γ-ATP, and referenced to the sum of all metabolites (tP) to investigate the stability of ATP across the experiment. Intracellular pH values were calculated from the chemical shift difference between Pi and PCr and [Mg2+] was calculated from the chemical shift difference between β-ATP and PCr ^1,8^.* |
| 5. Data Quality |  |
| a. Reported variables  (SNR, Linewidth (with reference peaks)) | *SNR was calculated using LCModel. Linewidth of the PCr peak is reported.* |
| b. Data exclusion criteria | *Linewidth of PCr peak > 15 Hz* |
| c. Quality measures of postprocessing Model fitting (e.g. CRLB, goodness of fit, SD of residual) |  |
| d. Sample Spectrum | *Figure 2* |

#### Marchenko-Pastur principal component analysis (MP-PCA) denoising

All denoising steps were performed in MATLAB (version R2024a, The Mathworks, Inc., USA). The raw data individual spectra were first phase- and frequency corrected. The resulting complex-valued FIDs were split into real and imaginary parts and organized into a matrix where the second dimension contained the time domain sampling and the first dimension a concatenation of all averages. More information about the theory behind the denoising can be found in Mosso et al. ^10^ . The resulting spectra were saved in raw format and quantified in the same way as described in the main manuscript.

#### Control Parameters for LCModel fitting

$LCMODL

xstep= 2

sddegz= 2

sddegp= 0.5

rfwhm= 3

ppmst= 8

ppmshf= -0.5;

ppmref(1,2)= -2.5

ppmend= -19

ppmcen= 0

owner= 'Center for Biomedical Imaging, Lausanne'

nunfil= 2048

nsimul= 0

nrefpk(2)= 1

neach= 999

nuse1= 5

namrel= 'g-ATP'

lps= 8

lprint= 6

lcoord= 9

key= xxxxxxx

hzpppm= 1.2031e+02

fwhmba= 0.049

TITLE = 'xxx'

FILPS = 'xxx.PS'

FILCOO = 'xxx.COORD'

FILRAW = 'xxx.RAW'

FILPRI = 'xxx.PRINT'

FILBAS= 'xxx.BASIS'

echot= 0.35

dkntmn= 2

desdt2= 10

desdsh= 0.01

deltat= 1.666e-04

degzer= 0

degppm= 0

deext2= 10

conrel= 3

chuse1= 'PCr','Pi','a-ATP','b-ATP','g-ATP'

ncombi=1

chcomb = 'NADH+NAD'

alpbst= 1.2e-9

alpbpn= 9.8e-10

alpbmx= 3.9e-7

alpbmn= 7.8e-10

$END

#### LCModel fit output example for one representative participant:

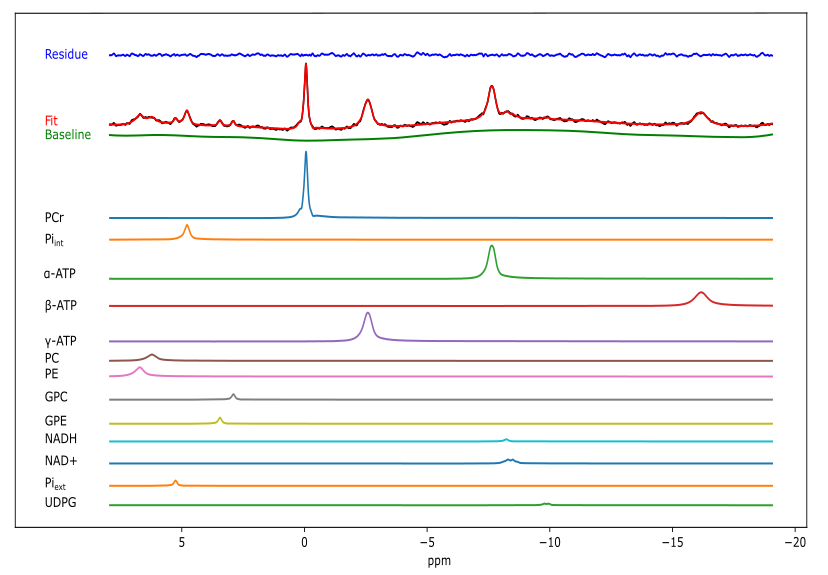

+

**Supplementary Figure 1:** LCModel fit output, including the separate metabolite functions.

### Example Overlap Plots of the fMRS voxel and fMRI BOLD maps:

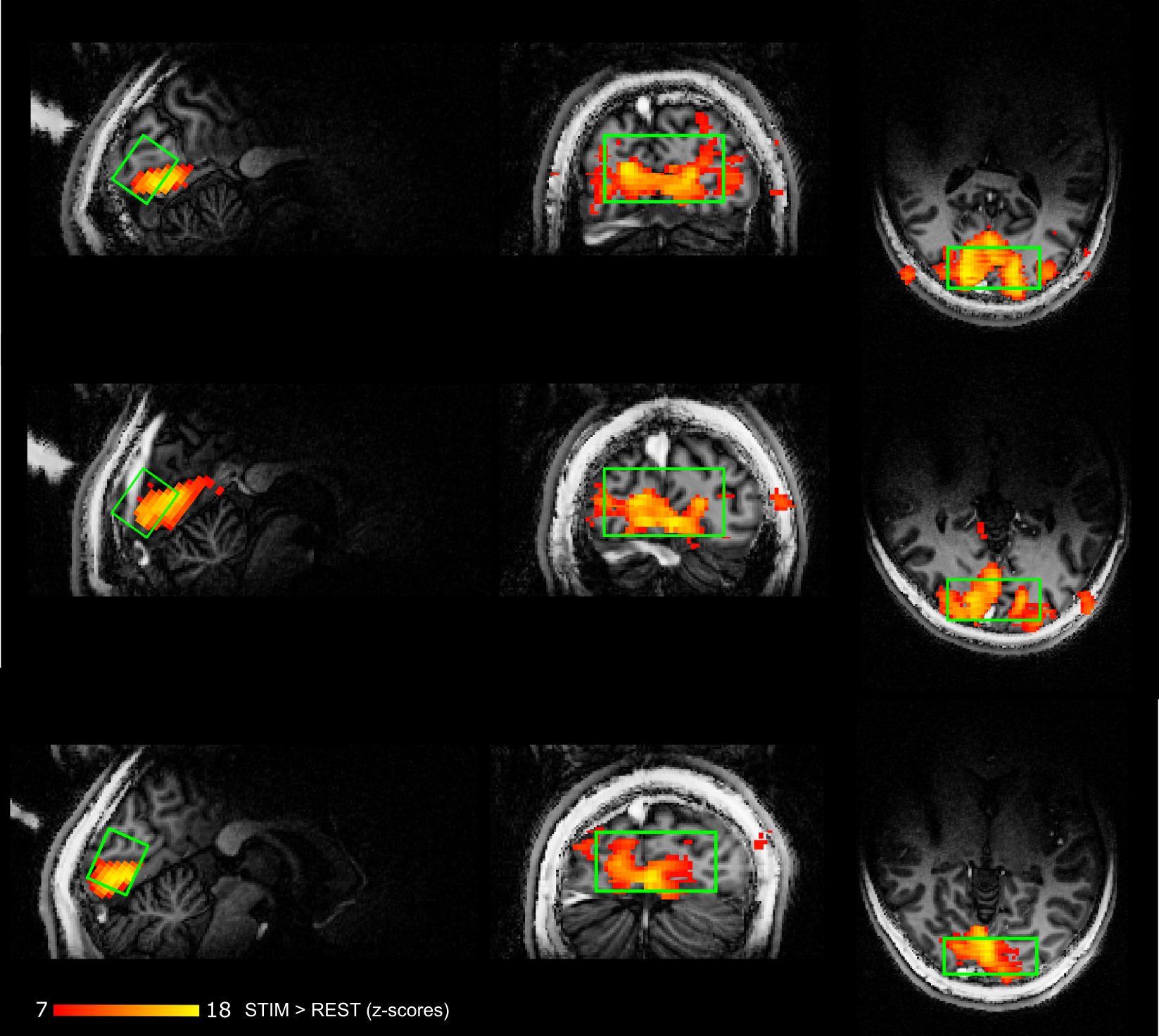

**Supplementary Figure 2:** Representative BOLD maps of three participants, with the fMRS voxel as overlay in green (Colorbar represents z-scores).

### UDPG one doublet vs. doublet of doublets fitting rationale

Using doublet of doublets to model UDPG introduces an additional degree of freedom on the quantification of NADH. Whether UDPG is detected or not influences the modelled NADH signal intensity. This increases uncertainty especially at low SNR conditions of dynamic ^31^P MRS and results in greater inter-participant variability in NADH estimates. Conversely, using one doublet UDPG modelling consistently leads to the expected slight overestimation (hence the NADH+ notation), but importantly, producing lower variance across subjects. It also improves the detection of UDPG in low SNR conditions as shown below.

– Number of participants in whom UDPG could be fit using the **doublet of doublets model**:

Data-points per block: REST1=11 (44%), STIM1=9 (36%), REST2=10 (40%), STIM2=8 (32%), REST3=10 (40%).

 – Mean ± **SD** of NADH (with explicit UDPG modelling): 0.09 ± **0.021**

 – Number of participants in whom UDPG could be fit using the **one doublet model**:

Data-points per block: REST1=14 (56%), STIM1=12 (48%), REST2=11 (44%), STIM2=10 (40%), REST3=11 (44%).

 – Mean ± **SD** of NADH+ (single doublet model): 0.10 ± **0.016**

These results confirm that NADH+ values are systematically larger, as expected due to inclusion of UDPG signal at -8.23 ppm, and exhibit lower inter-participant variability. In addition, the single-doublet model yielded a higher number of successful UDPG detections per block, indicating improved fitting robustness. Together, the reduced variance and increased number of successfully fitted data points support the approach that the one-doublet modelling strategy provides more stable estimates for assessing temporal dynamics of UDPG and NADH+. Furthermore, since we did not observe a temporal difference in UDPG, it is unlikely that UDPG contributes to NADH+ dynamics.

### Supplementary Results

#### Without MP-PCA applied:

|  | Estimated mean parameters [95% CIs] | | | | |  |  |
| --- | --- | --- | --- | --- | --- | --- | --- |
| Metabolite | Rest 1 | Stim 1 | Rest 2 | Stim 2 | Rest 3 | Statistics  (block+linewidth) | Post-Hoc test |
| Linewidth (Hz) | 12.1 [11.7 12.5] | 12.2 [11.8 12.6] | 12.6 [12.2 13.00] | 12.5 [12.1 12.8] | 12.5 [12.1 12.8] | F(4,88) = 1.69, p = 0.15 | - |
| PCr (mM) | 2.84 [2.63 3.06] | 2.81 [2.59 3.02] | 2.92 [2.71 3.13] | 2.85 [2.64 3.06] | 3.03 [2.82 3.24] | F(4,88) = 0.79, p = 0.53 | - |
| Pi_int_ (mM) | 0.81 [0.70 0.92] | 0.88 [0.78 0.99] | 0.81 [0.70 0.92] | 0.82 [0.71 0.93] | 0.77 [0.67 0.88] | F(4,88) = 0.59, p = 0.67 | - |
| PCr/Pi_int_ (-) | 3.71 [3.27 4.13] | 3.35 [2.92 3.77] | 3.79 [3.37 4.21] | 3.59 [3.17 4.02] | 4.1 [3.66 4.53] | F(4,86) = 1.62, p = 0.17 | - |
| NAD^+^ (mM) | 0.23 [0.18 0.28] | 0.16 [0.12 0.21] | 0.25 [0.21 0.29] | 0.29 [0.19 0.29] | 0.19 [0.15 0.24] | **F(4,80) = 2.56, p=0.04** | **REST1>STIM1 t(80) = 2.9, p=0.03;**  **STIM1<REST2 t(80) = -2.6, p=0.04** |
| NADH (mM) | 0.11 [0.05 0.18] | 0.12 [0.06 0.17] | 0.08 [0.04 0.14] | 0.09 [0.04 0.14] | 0.08 [0.04 0.16] | F(4,47) = 0.33, p = 0.85 | - |
| tNAD (mM) | 0.26 [0.19 0.32] | 0.23 [0.16 0.29] | 0.32 [0.25 0.38] | 0.30 [0.24 0.37] | 0.24 [0.18 0.30] | F(4,81) = 1.62, p = 0.18 | - |
| pH_int_ | 6.987 [6.982 6.992] | 6.989 [6.985 6.995] | 6.989 [6.984 6.994] | 6.993 [6.988 6.998] | 6.994 [6.989 6.999] | F(4,88) = 1.24, p = 0.30 | - |
| pH_ext_ | 7.378 [7.376 7.381] | 7.378 [7.375 7.381] | 7.377 [7.373 7.379] | 7.376 [7.373 7.378] | 7.376 [7.373 7.379] | F(4,88) = 1.02, p = 0.40 | - |
| [Mg^2+^] (mM) | 0.175 [0.172 0.178] | 0.177 [0.174 0.179] | 0.174 [0.171 0.176] | 0.174 [0.170 0.176] | 0.174 [0.171 0.177] | F(4,88) = 1.27, p = 0.29 | - |
| ATP/tP ( -) | 0.201 [0.194 0.209] | 0.196 [0.190 0.205] | 0.195 [0.188 0.203] | 0.194 [0.186 0.201] | 0.199 [0.192 0.207] | F(4,88) = 0.82, p = 0.51 | - |
| UDPG (mM) | 0.32 [0.24 0.41] | 0.28 [0.18 0.38] | 0.29 [0.19 0.38] | 0.31 [0.21 0.42] | 0.26 [0.17 0.36] | F(4,38) = 0.32, p = 0.86 | - |

**Supplementary Table 2:** **Statistical analysis for the fitting using the doublet of doublets modelling.** The results of the main analysis and post-hoc tests conducted in our study are shown. The analysis was performed using linear mixed effects models to estimate the means and confidence intervals for each parameter. Post-hoc comparisons were conducted using Tukey's HSD tests. The parameters investigated include linewidth and various metabolites and physiological parameters. The estimated means are presented alongside their 95% confidence intervals (CI) per block and associated statistical significance values (p-values). Concentrations of NAD⁺, NADH, tNAD, UDPG are given in mM, referenced to an assumed ꭤ-ATP concentration of 2.8 mM, PCr, Piint, are given in mM, referenced to an assumed γ-ATP concentration of 2.8 mM in the human occipital lobe. [Mg²⁺] is reported in mM. ATP was calculated as an average of α-ATP, β-ATP and γ-ATP, and referenced to the sum of all metabolites (tP).

|  | Estimated mean % CRLB  [95% CIs] | Estimated mean absolute CRLB  [95% CIs] |
| --- | --- | --- |
| Metabolite |  |  |
| PCr | 2.96 [2.65 3.26] | 0.08 [0.07 0.09] |
| Pi_int_ | 10.63 [9.19 12.06] | 0.09 [0.07 0.11] |
| Pi_ext_ | 28.68 [22.10 35.26] | 0.011 [0.010 0.012] |
| NAD^+^ | 19.77 [16.02 23.53] | 0.05 [0.04 0.07] |
| NADH+ | 90.94 [8.76 173.12] | 0.08 [0.007 0.15] |
| UDPG | 26.43 [20.43 32.43] | 0.014 [0.013 0.014] |

**Supplementary Table 3: CRLB.** The average Cramer Rao Lower Bound (CRLB) per fitted metabolite over all blocks and all participants is shown here (estimated mean and 95% confidence interval (CI)) in percentage and as concentration values ^9^.

##### Pi_ext_ quantification details:

Data-points per block: REST1=8 (32%), STIM1=13 (52%), REST2=10 (40%), STIM2=6 (24%), REST3=11 (44%).

##### NADH+ quantification details:

Data-points per block: REST1=12 (48%), STIM1=14 (56%), REST2=16 (64%), STIM2=18 (72%), REST3=15 (60%).

##### UDPG concentration changes:

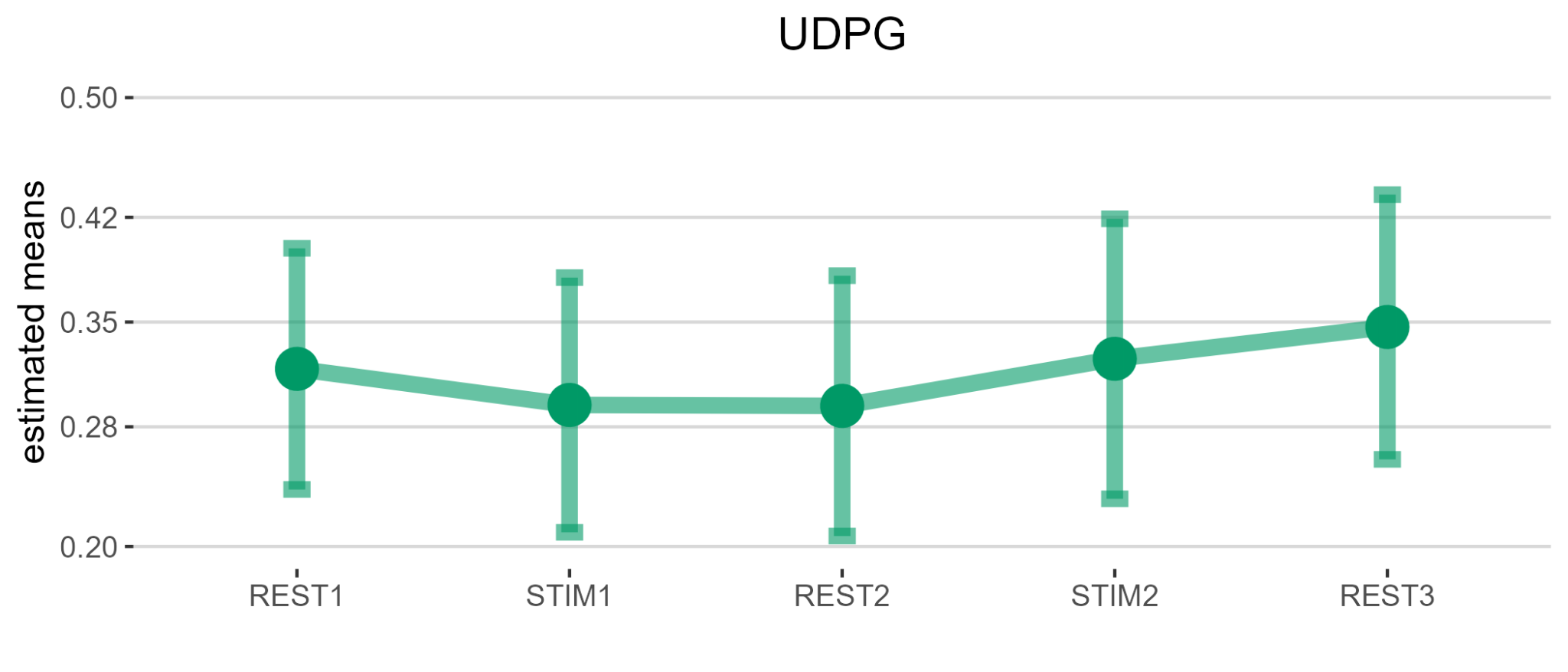

**Supplementary Figure 3:** UDPG concentration changes over the 5 blocks.

UDPG data-points per block: REST1=14 (56%), STIM1=12 (48%), REST2=11 (44%), STIM2=10 (40%), REST3=11 (44%).

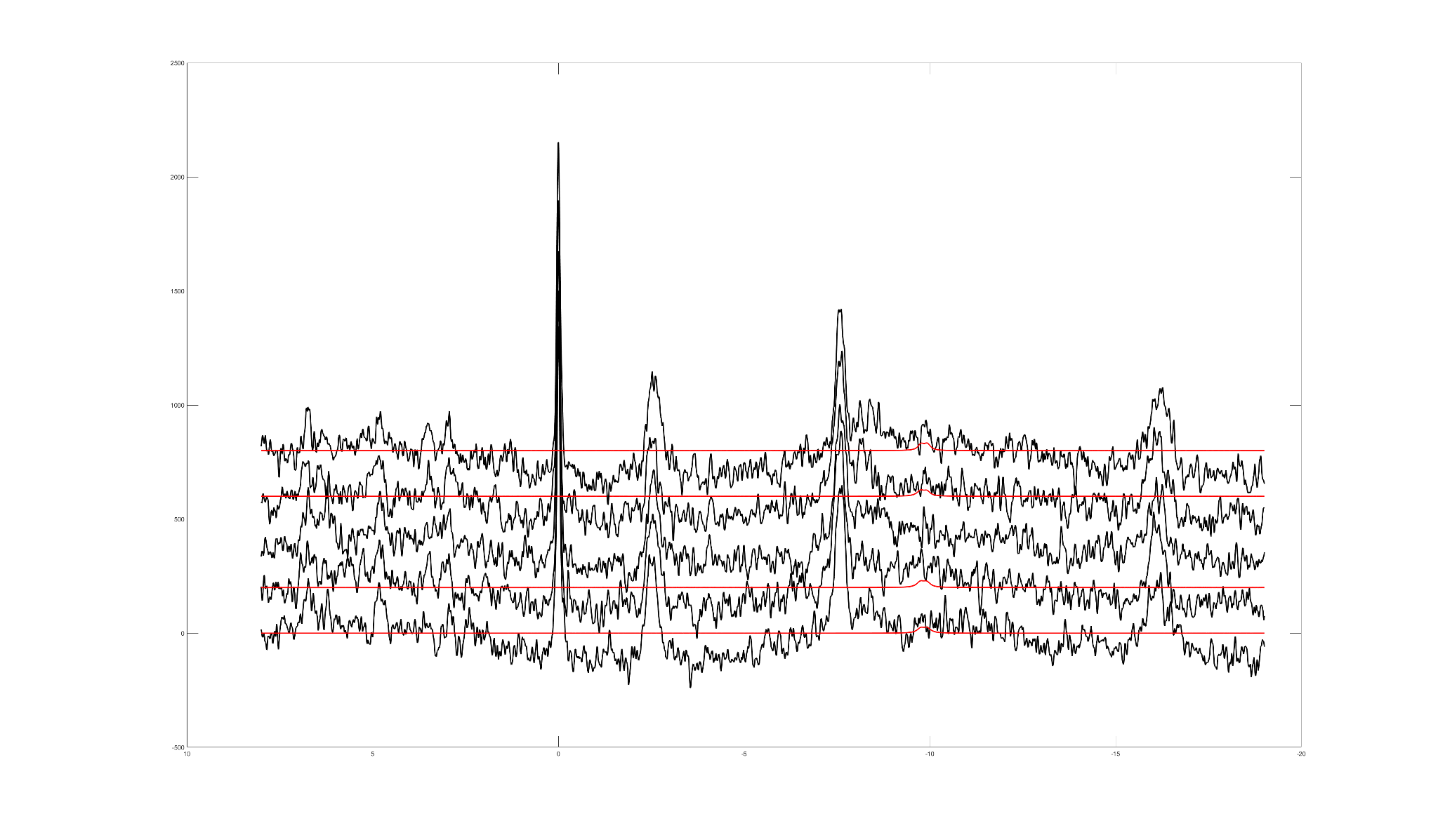

**Supplementary Figure 4:** Example fit of UDPG (in red) per block (REST1, STIM1, REST2, STIM2, REST3 from bottom to top) of one representative participant. UDPG could not be reliably fitted in REST2 and is therefore not shown.

#### Additional BOLD effect check:

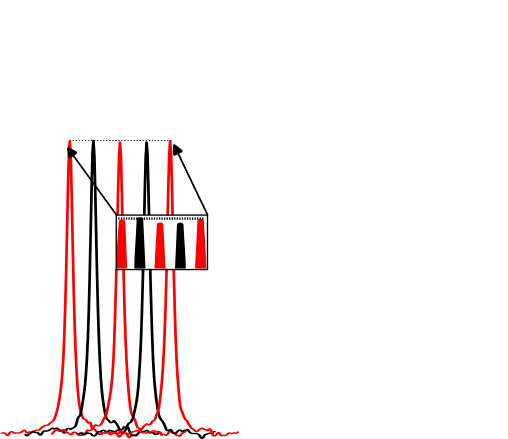

**Supplementary Figure 5:** Peak height of PCr (averaged over all participants) over the 5 blocks of the experiment (with a zoomed in highlight box for the peaks), no significant changes are visible. Black=rest, red=activation.

#### Pi_int_ and Pi_ext_ fit

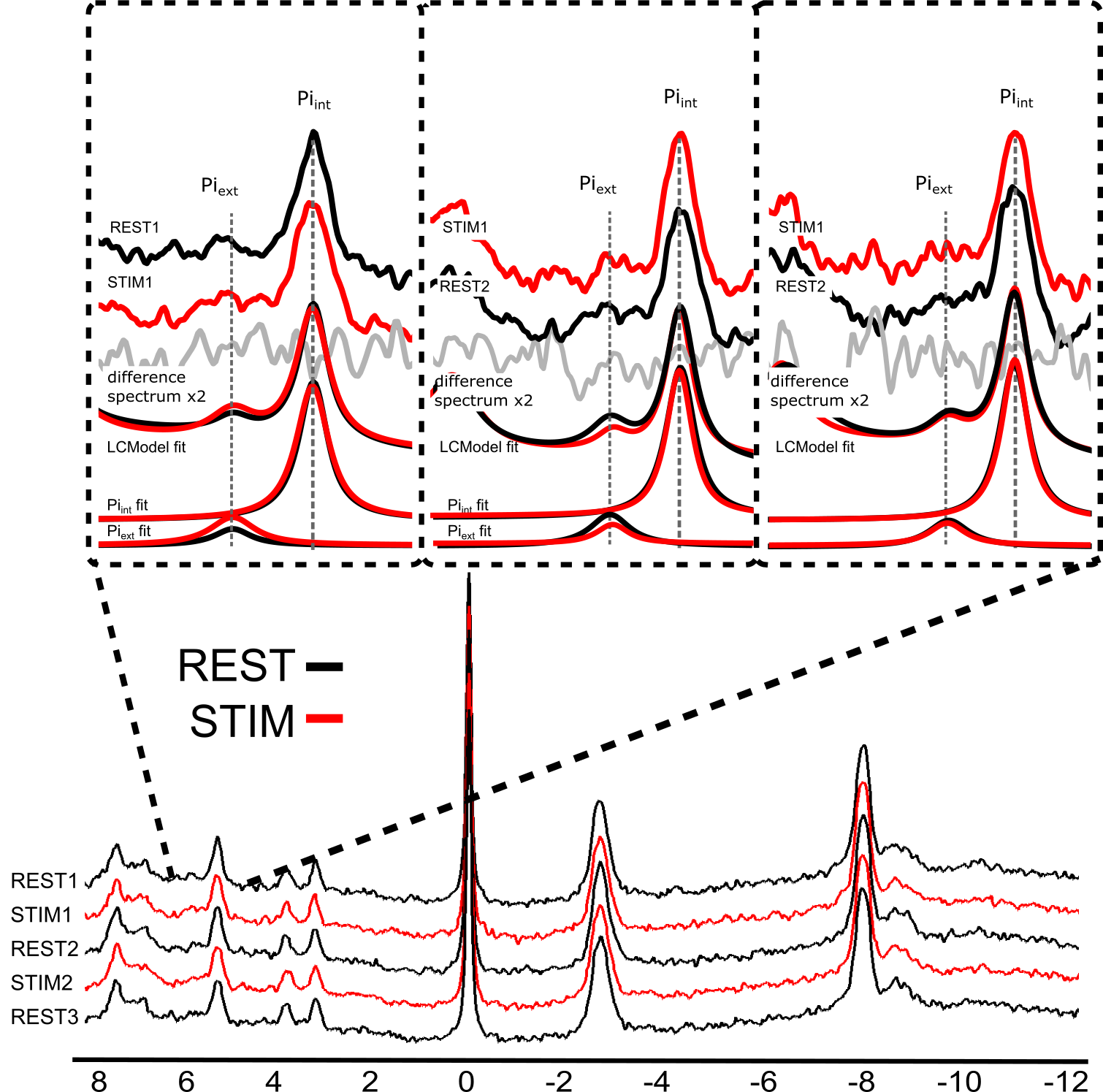

**Supplementary Figure 6:** Quality measures: All spectra of all included participants during the rest (black) and activation (red) blocks were averaged together and plotted above each other, with a highlight on the Pi area around 5 ppm, including a difference-spectrum of rest-activation (gray), and the Pi_int_ and Pi_ext_ fits by LCModel.

#### 10-block analysis:

| Metabolite | Statistics (block+linewidth)  Over all 10 blocks | Post-Hoc test |
| --- | --- | --- |
| Linewidth (Hz) | F(9,198)=0.68, p=0.73 | - |
| PCr (mM) | F(9,194)=0.77, p=0.64 | - |
| Pi_int_(mM) | F(9,193)=1.38, p=0.20 | - |
| PCr/Pi (-) | F(9,192)=1.53, p=0.13 | - |
| NAD^+^ (mM) | F(9,164)=1.76, p=0.07 | - |
| NADH+ (mM) | F(9,111)=1.11, p=0.36 | - |
| tNAD (mM) | F(9,189)=1.74, p=0.08 | - |
| pH_int_ | F(9,197)=0.73, p=0.68 | - |
| pH_ext_ | F(9,197)=0.51, p=0.87 | - |
| [Mg^2+^] (mM) | F(9,197)=0.21, p=0.99 | - |
| ATP/tP (-) | F(9,197)=0.75, p=0.66 | - |
| UDPG (mM) | F(9,83)=1.47, p = 0.61 | - |

**Supplementary Table 4:** The results of the exploratory analysis, dividing the blocks into two halves, and post-hoc tests are presented. The analysis was performed using linear mixed effects models to estimate the means and confidence intervals for each parameter. Post-hoc comparisons were conducted using Tukey's HSD tests. The parameters investigated include linewidth and various metabolites and physiological parameters. The statistical significance values (p-values) are shown for the effect of timepoint (‘’block’’).

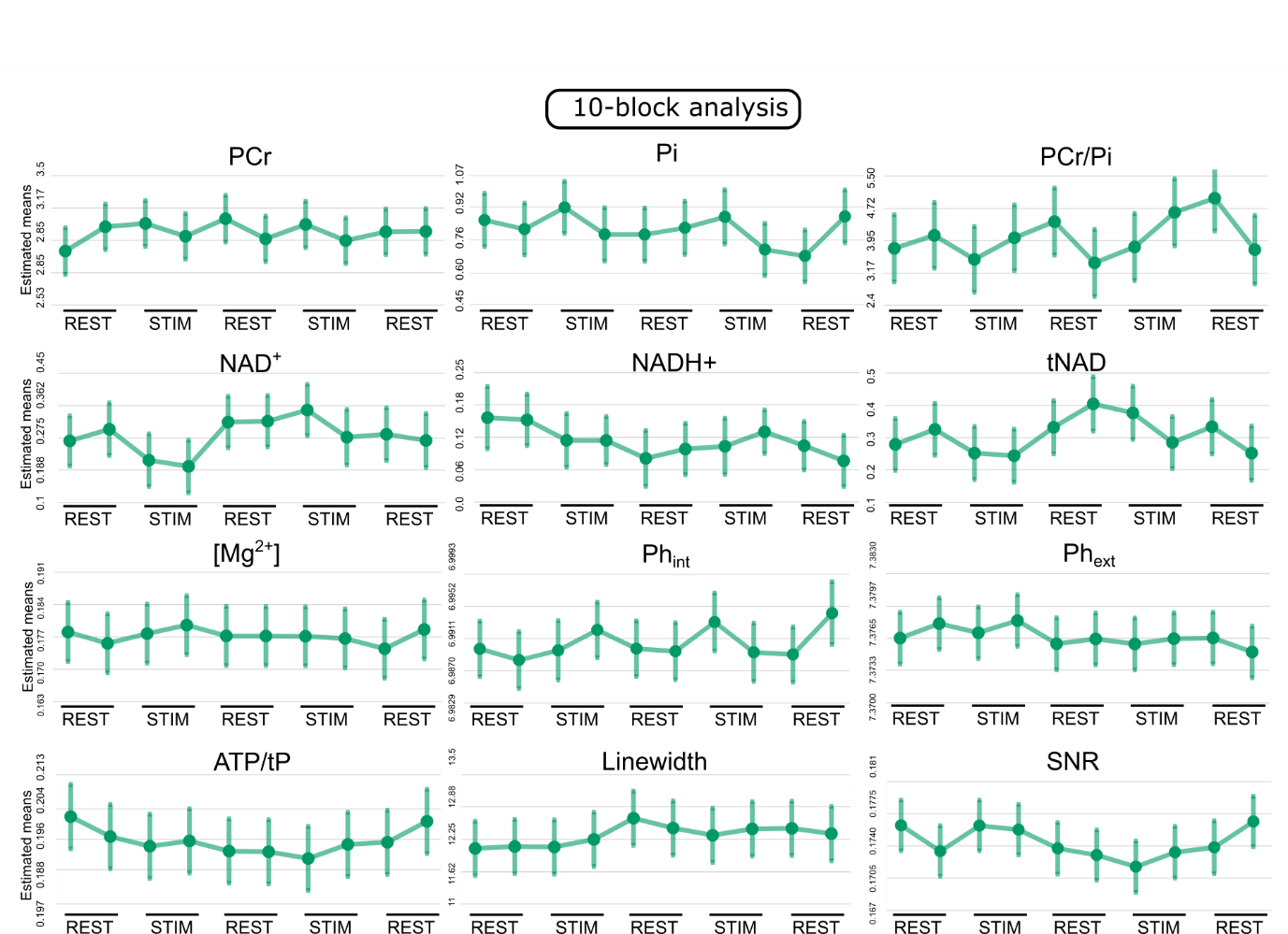
**Supplementary Figure 7: Results.** Exploratory analysis for individual concentration changes per 1/2 block (two concentration averages per rest1, stim1, rest2, stim2, rest3) for PCr, Pi, PCr/Pi, NAD^+^,NADH+. tNAD, [Mg^2+^], pH_int_, pH_ext_, ATP/tP, Linewidth, and signal-to-noise-ratio.

#### Results for UDPG:

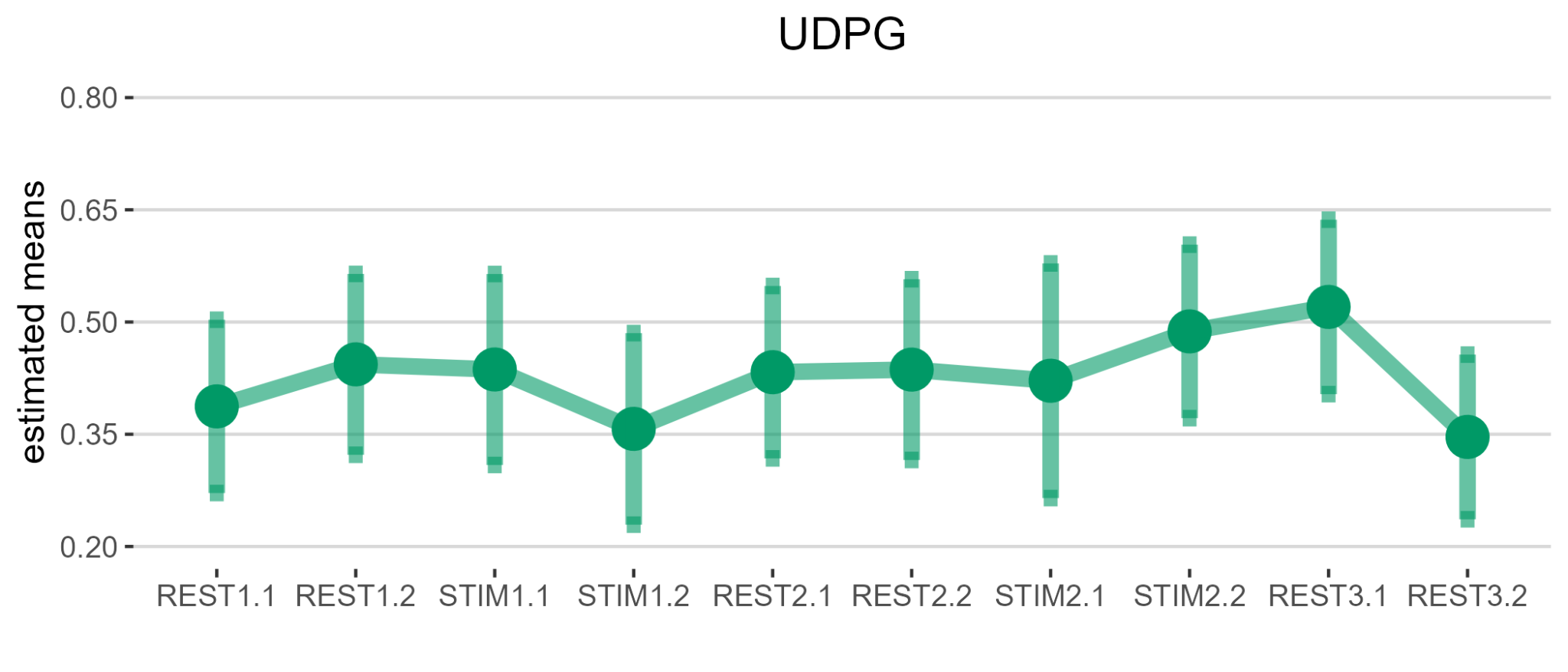

**Supplementary Figure 8:** UDPG concentration changes over the 10 half-blocks. Around 40% of the datapoints were quantifiable.

#### With MP-PCA applied:

|  | Estimated mean parameters [95% CIs] |  |  |
| --- | --- | --- | --- |
| Metabolite | Mean over all blocks | Statistics (block+linewidth) | Post-Hoc test |
| Linewidth (Hz) | 6.86 [6.07 7.65] | F(4,96)=1.14, p=0.34 | - |
| PCr (mM) | 3.03 [2.78 3.28] | F(4,78)=0.63, p=0.64 | - |
| Pi_int_ (mM) | 0.81 [0.74 0.89] | F(4,73)=1.78, p=0.14 | - |
| PCr/Pi (-) | 3.70 [3.36 4.05] | F(4,73)=1.59, p=0.19 | - |
| NAD^+^ (mM) | 0.26 [0.22 0.30] | **F(4,75)=3.23, p=0.02** | **rest1>stim1: t(74)=2.80, p=0.04;**  **stim1<rest2: t(76)=-2.95, p=0.03** |
| NADH+ (mM) | 0.04 [0.03 0.05] | F(4,109)=0.67, p=0.61 | - |
| tNAD (mM) | 0.31 [0.25 0.36] | **F(4,67)=2.79, p=0.03** | rest1>stim1: t(67)=2.66, p=0.07;  **stim1<rest2: t(66)=-3.09, p=0.02** |
| pH_int_ | 6.996 [6.994 6.998] | F(4,78)=0.26, p=0.90 | - |
| [Mg^2+^] (mM) | 0.166 [0.165 0.168] | F(4,70)=0.83, p=0.51 | - |
| ATP/tP (-) | 0.208 [0.203 0.212] | F(4,58)=1.47, p=0.22 | - |
| UDPG (mM) | 0.47 [0.36 0.59] | F(4,84)= 0.81, p=0.61 | - |

**Supplementary Table 5: Statistical analysis:** The results of the main analysis and post-hoc tests conducted on the denoised data. The analysis was performed using linear mixed effects models to estimate the means and confidence intervals for each parameter. Post-hoc comparisons were conducted using Tukey's HSD tests. The parameters investigated include linewidth and various metabolites and physiological parameters. The estimated means are presented alongside their 95% confidence intervals (CI) and associated statistical significance values (p-values).

**Supplementary Figure 9: Results of the data with MP-PCA denoising applied.** A) Quality measures: Linewidth of the PCr peak over the course of the experiment. Additionally, all spectra during the rest (black) and activation (red) blocks were averaged together and plotted above each other, with a highlight on the NAD^+^ area around -9ppm, including a difference-spectrum of rest-activation (gray). B) Individual concentration changes per block (5-block analysis; rest1, stim1, rest2, stim2, rest3) for PCr, Pi, PCr/Pi, NAD^+^, NADH+, tNAD, ATP/tP, pH, and [Mg2+]. *=p<0.02

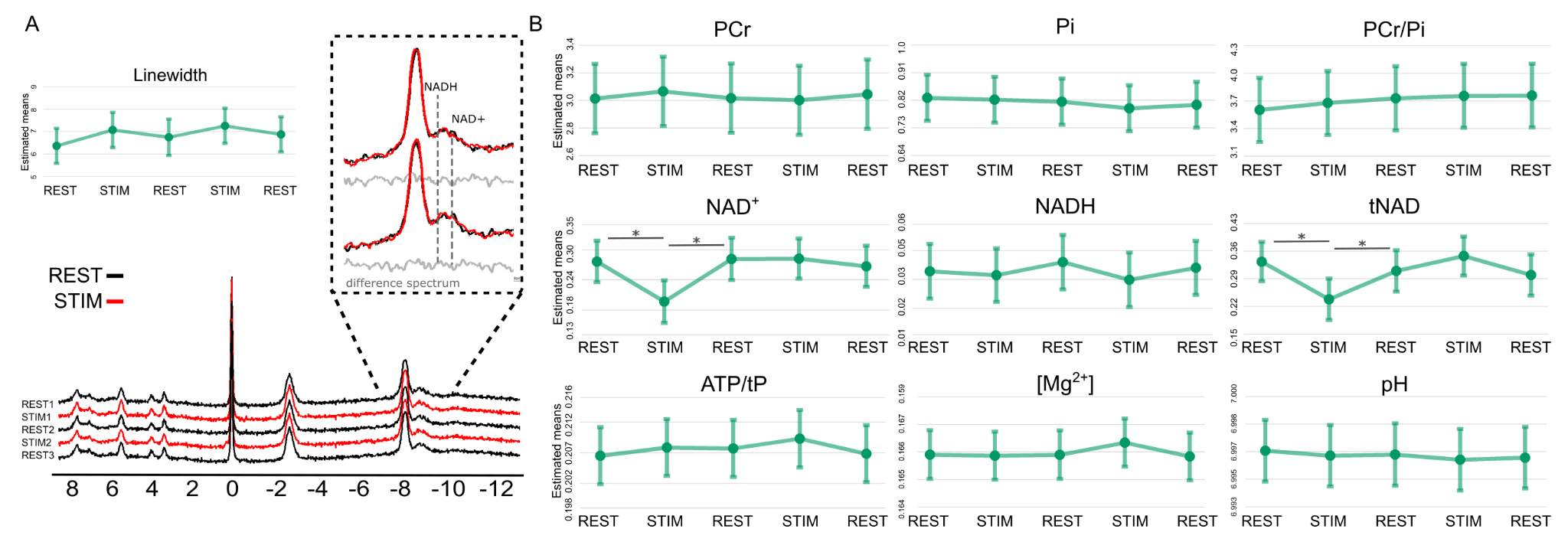

+

+
